# NorA, HmpX, and NorB cooperate to reduce NO toxicity during denitrification and plant pathogenesis in *Ralstonia solanacearum*

**DOI:** 10.1101/2021.11.08.467854

**Authors:** Alicia N. Truchon, Connor G. Hendrich, Adam F. Bigott, Beth L. Dalsing, Caitilyn Allen

## Abstract

*Ralstonia solanacearum*, which causes bacterial wilt disease of many crops, needs denitrifying respiration to succeed inside its plant host. In the hypoxic environment of plant xylem vessels this pathogen confronts toxic oxidative radicals like nitric oxide (NO), which is generated by both bacterial denitrification and host defenses. *R. solanacearum* has multiple distinct mechanisms that could mitigate this stress, including Repair of Iron Cluster (RIC) homolog NorA, nitric oxide reductase NorB, and flavohaemoglobin HmpX. During denitrification and tomato pathogenesis and in response to exogenous NO, *R. solanacearum* upregulated *norA, norB*, and *hmpX*. Single mutants lacking *ΔnorB, ΔnorA*, or *ΔhmpX* increased expression of many iron and sulfur metabolism genes, suggesting that losing even one NO detoxification system demands metabolic compensation. Single mutants suffered only moderate fitness reductions in host plants, possibly because they upregulated their remaining detoxification genes. However, *ΔnorA/norB, ΔnorB/hmpX*, and *ΔnorA/hmpX* double mutants grew poorly in denitrifying culture and *in planta*. Loss of *norA, norB*, and *hmpX* may be lethal, since the methods used to construct the double mutants did not generate a triple mutant. Aconitase activity assays showed that NorA, HmpX and especially NorB are important for maintaining iron-sulfur cluster proteins. Additionally, plant defense genes were upregulated in tomatoes infected with the NO-overproducing *ΔnorB* mutant, suggesting that bacterial detoxification of NO reduces pathogen visibility. Thus, *R. solanacearum*’s three NO detoxification systems each contribute to and are collectively essential for overcoming metabolic oxidative stress during denitrification, for virulence and growth in tomato, and for evading host plant defenses.

**Importance:** The soilborne plant pathogen *Ralstonia solanacearum* (*Rs*) causes bacterial wilt, a serious and widespread threat to global food security. *Rs* is metabolically adapted to low oxygen conditions, using denitrifying respiration to survive in the host and cause disease. However, bacterial denitrification and host defenses generate nitric oxide (NO), which is toxic and also alters signaling pathways in both plants and the pathogen. *Rs* mitigates NO with a trio of mechanistically distinct proteins: NO-reductase NorB, Repair of Iron Centers NorA, and oxidoreductase HmpX. This redundancy, together with analysis of mutants and *in-planta* dual transcriptomes, indicates that maintaining low NO levels is integral to *Rs* fitness in tomatoes (because NO damages iron-cluster proteins) and to evading host recognition (because bacterially produced NO can trigger plant defenses).

## Introduction

*Ralstonia solanacearum* (*Rs*), a soil-dwelling plant pathogen, causes bacterial wilt disease on a wide range of economically important plants, including tomatoes. Bacterial wilt is a serious socioeconomic problem in tropical regions, especially in developing countries where crop loss can be devastating for subsistence farmers (1). To date, there is no effective control strategy to combat bacterial wilt (2). *Rs* draws on its broad repertoire of metabolic capabilities to survive in soil and water, invade plant roots, and colonize and obstruct its host’s water-transporting xylem vessels (3). The pathogen’s metabolism adapts rapidly as it transitions among diverse micro-niches in surface water, soil, and inside hosts (4, 5). Plant xylem vessels, the primary in-host habitat of *Rs*, contain little oxygen but have substantial levels of nitrate (NO_3_), around 30 mM (6).

Bacteria have several ways to make ATP under low oxygen conditions. These include fermentation and respiration using alternate terminal electron acceptors (TEAs) like sulfur, iron, and nitrogen (7). Nitrate respiration and denitrification require a series of membrane-bound and periplasmic enzymes that reduce NO_3_^-^ stepwise to dinitrogen gas (N_2_) (8). Denitrifying respiration allows organisms to produce energy from NO_3_^-^ in hypoxic environments such as soil, marine sediments, landfills, wastewater treatment plants, bioreactors, and inside eukaryotic hosts (8–15).

Nitrate metabolism is broadly conserved across plant pathogenic *Ralstonia spp*. (16–18). *Rs* strain GMI1000 has a complete pathway for denitrifying respiration wherein the nitrate reductase NarG reduces NO_3_^-^ to NO_2_^-^ (nitrite); the nitrite reductase AniA reduces NO_2_^-^ to NO (nitric oxide); the nitric oxide reductase NorB converts NO to N_2_O (nitrous oxide); and finally, the nitrous oxide reductase NosZ converts N_2_O to N_2_ (Fig. 1). When *Rs* invades tomato stems, xylem oxygen levels decline even further and the pathogen’s denitrification genes are substantially upregulated (6, 19). We previously established that *Rs* uses NO_3_^-^ and its reduction products as TEAs to generate proton motive force that drives ATP synthesis (6). Possibly as a result, denitrification contributes quantitatively to *Rs* growth *in planta* and to bacterial wilt virulence (6, 19).

**Figure 1.**
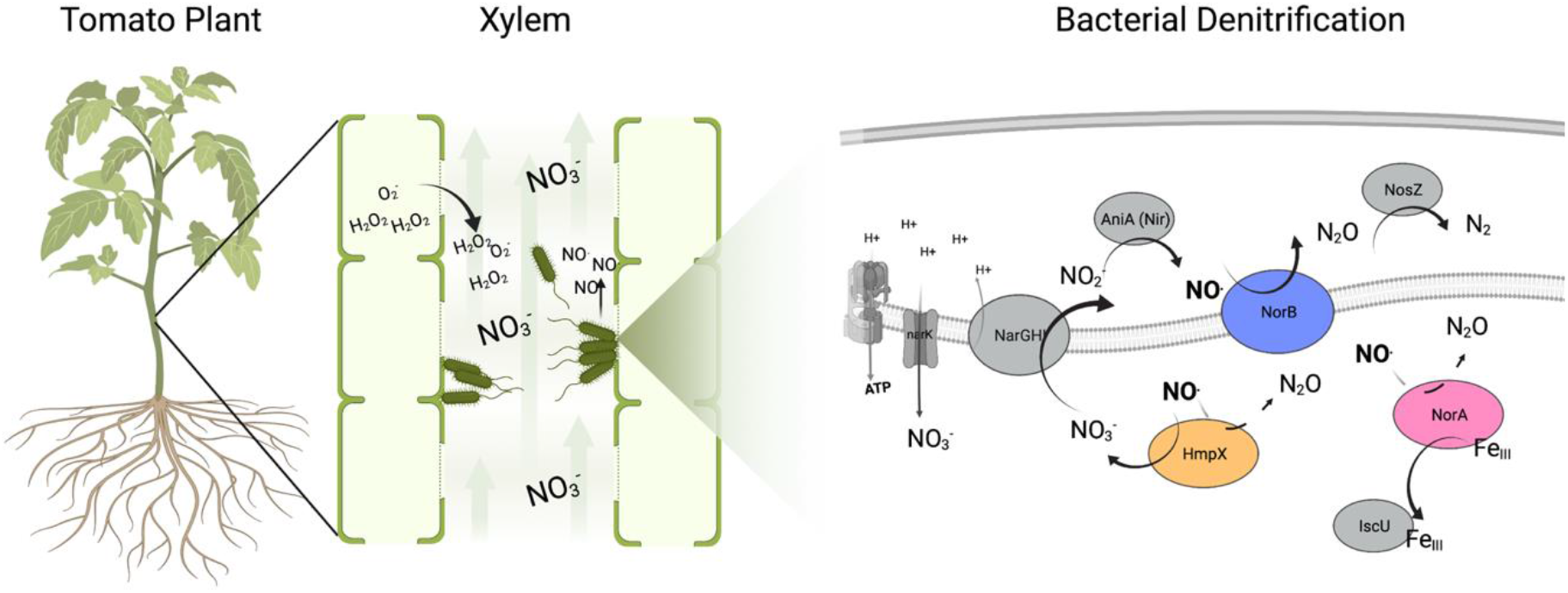
*Ralstonia solanacearum* strain GMI1000 denitrifies in tomato xylem, generating energy and NO. Tomato plant xylem contains ~30 mM NO_3_^-^. *Rs* uses denitrifying respiration to reduce NO_3_^-^ and generate ATP. Denitrification reduces NO_3_^-^ to N_2_ gas via the enzymes NarGHI, AniA, NorB, and NosZ. NO_3_^-^ reduction generates toxic NO that must be oxidized, reduced, or sequestered to prevent cellular damage. In response to bacterial infection, tomato plants also produce oxidative compounds, including H_2_O_2_, and O_2_^-^. *R. solanacearum* NorA, NorB, and HmpX proteins interact with NO by either reducing NO to N_2_O, oxidizing NO to NO_3_^-^, or repairing iron centers damaged by NO. Figure created in part using BioRender.

However, denitrifying respiration comes at a cost. The pathway generates two highly reactive nitrogen species (RNS): NO_2_^-^ and NO (8, 15). NO_2_^-^ and NO are toxic in their own right because they damage iron centers of important iron-sulfur cluster (Fe-S) heme proteins (20, 21). NO is both a RNS and a reactive oxygen species (ROS) that interacts with other oxygen or nitrogen species to form even more damaging species like peroxynitrite (22). These secondary NO products have highly toxic effects on major cellular components including metalloproteins, lipids, and nucleic acids (22, 23).

In addition to the NO generated by prokaryotic respiration, microbes encounter NO produced by their eukaryotic hosts (24, 25). NO can act as diffusible signal that does not require a carrier, and is a major plant signaling molecule that rapidly regulates plant defense functions, including cell death (26, 27). Many hosts also produce NO and H_2_O_2_ to directly kill pathogens (28–31). In response to this oxidative attack, animal and plant pathogens including *Erwinia spp., Pseudomonas spp., Staphylococcus aureus*, and *Neisseria gonorroheae* use denitrification pathway enzymes like NO_2_^-^ reductase (NIR) and NO reductase (NOR) not only to produce energy but also to reduce the toxic load of RNS, sometimes by decoupling them from the electron transport chain (8, 12). Microbes have evolved additional specialized mechanisms to mitigate RNS stress (6, 8, 29, 30, 32–36). The flavohaemoglobin Hmp is an oxidoreductase that uses a globin-like NO-binding domain, NAD, and FAD to catalyze the conversion of NO and O_2_ to NO_3_^-^ when oxygen is available, or to reduce NO to N_2_O in the absence of O_2_ (28, 30, 37, 38). Homologs of Hmp are present across the bacterial domain (28, 29, 39). A second protective mechanism involves NO-inducible Repair of Iron Centers (RIC) di-iron proteins. RICs use a hemerythrin-like domain to decrease oxidative damage by interacting with iron storage proteins like Dps and IscU to deliver iron to damaged Fe-S clusters (36, 40–44).

The *Rs* GMI1000 genome has genes encoding a putative RIC protein, NorA; a NO reductase, NorB; and an oxidoreductase, HmpX. When *Rs* grows in tomato xylem, *norA, norB*, and *hmpX* are upregulated 75-, 51-, and 43-fold respectively, relative to when *Rs* grows in rich media (19). These were among the most differentially expressed genes *in planta*, where *Rs* cells experience an oxidative environment (19, 31, 45). This upregulation implied that during plant pathogenesis, *Rs* depends on the products of *norA, norB*, and *hmpX* to mitigate oxidative stress produced by its own denitrifying respiration and by the plant host. This functional redundancy suggested that detoxifying NO is critically important for *R. solanacearum*. We tested this hypothesis using a panel of single and double mutants lacking *norA, hmpX*, and *norB* combined with transcriptomic, biochemical, and plant assays.

## Materials and Methods

### Bacterial Growth Conditions

The *R. solanacearum* and *Escherichia coli* strains used are listed in Table S5. *E. coli* strains were grown in LB broth and *R. solanacearum* strains were grown on rich CPG media at 28°C, shaking at 225 rpm unless otherwise noted. As appropriate, antibiotics were used at the following concentrations: 25 ug/ml kanamycin and 10 ug/ml tetracycline. We grew bacteria under the previously determined denitrifying conditions: in VDM media modified with 30 mM NO_3_ in low oxygen (either 0 or 0.1% O_2_), with shaking at 225 rpm or on medium speed in a microplate reader (BioTek, Winooski, VT, USA) (6). For hypoxic assays, bacteria were grown in an anaerobic chamber (InvivO2, Baker Ruskinn, Sanford, Maine, USA) set to 0.1% O_2_. For anoxic assays, cells were grown in GasPak pouches (BD, Franklin Lakes, NJ, USA).

### Mutant construction

All *Rs* mutants were constructed in phylotype I sequevar 18 strain GMI1000. Unmarked Δ*norA* and Δ*norR* mutants lacking the complete *norA* or *norR* ORF were generated using Gibson assembly and *sacB* positive selection vector pUFR80 as described.(46) Briefly, PCR with KapaHifi DNA-polymerase was used to amplify up- and down-stream regions of Rsp0958 (*norA*) or Rsp0959 (*norR*); PCR fragments were annealed with pUFR80 to form either pUFR80-*norA* or pUFR80-*norR*, which were then transformed into GMI1000; and kanamycin and sucrose selection were used to generate clean in-frame deletion mutants. Double mutants were made by transforming previously constructed plasmids into the Δ*norA* or previously constructed Δ*norB* and Δ*hmpX* mutant backgrounds (6). All mutants were confirmed with sequencing. All primers and mutant strains are listed in Supporting Information Table S5.

### Plant experiments

Disease assays were conducted as previously described (47). Briefly, wilt-susceptible cv. Bonny Best tomato plants were grown at 28°C with a 12 h day-night light cycle and watered daily with 0.5X strength Hoagland’s solution. Two-week-old seedlings were transplanted into 4-inch pots containing ~80 g potting mix. Two days later, unwounded plants were inoculated by drenching the soil with 50 mL of a 1×10^8^ CFU/mL bacterial suspension. Inoculum was determined turbidometrically and confirmed by dilution plating as described (48). Plant wilt symptoms were rated using a 0-4 disease index for 14 days (48).

### Alignments

NCBI BLASTp non-redundant protein sequence database (https://blast.ncbi.nlm.nih.gov) was used to compare percent amino acid identity (% AA ID) and percent query cover (% QC) of *R. solanacearum* NorA, NorB, and HmpX to *C. necator, N. gonorrhoeae, S. aureus, E. coli, S. enterica*, and *X. fastidiosa*. MUSCLE multiple sequence alignment tool (https://www.ebi.ac.uk/Tools/msa/muscle) was used to align *R. solanacearum* NorA, NorB, and HmpX amino acid sequences with homologs in *C. necator, N. gonorrhoeae, S. aureus, E. coli, S. enterica*, and *X. fastidiosa*.

### RNA extraction and Transcriptomic Analyses

#### RNA extraction

RNA was collected from denitrifying *R. solanacearum* bacterial cultures or from stem tissue of plants 72 h after petiole-inoculation with *Rs* strain GMI1000, *ΔnorB*, or water.

For transcriptomes of cultured cells, bacteria were grown in triplicate in VDM +30 mM NO_3_^-^ without shaking for 16 h at 28°C in 0.1% O_2_. Sub-samples were dilution-plated to determine CFU/ml, then samples were centrifuged at room temperature for 5 minutes at 3000 x *g*, supernatant was removed, and pellets were frozen in liquid nitrogen. Total RNA was extracted using a modified version of the Quick-RNA™ MiniPrep Kit (Zymo Research, Irvine, CA, USA). Briefly, frozen pellets were resuspended in 400 μL cold TE pH 8 with 1 mg/mL lysozyme, 0.25 μL Superase inhibitor (Ambion, Austin, TX, USA), and 80 μL OF 10% SDS, vortexed for 10s, then transferred to a new 2 mL tube, shaken at 300 rpm for 2 min. 800 μL of RNA-Lysis buffer was added, then samples were cleaned according to the kit manufacturer’s instructions. Samples were eluted in 100 μL nuclease-free water then DNA was then removed using the DNA-free DNAse kit according to manufacturer’s instructions for Rigorous DNAse treatment (Invitrogen, Carlsbad, CA, USA). After DNAse inactivation, samples were further cleaned by chloroform extraction, precipitated overnight at −20°C with 100 μM Sodium Acetate pH 5.5 and 66% ethanol. Samples were checked for concentration on a Nanodrop (Thermo Fischer Scientific, Wilmington, DE, USA), for DNA contamination by PCR using the qRT-PCR primers *serC*_F/R, and for RNA integrity (RIN) using Agilent bioanalyzer 21000 (Agilent, Santa Clara, CA, USA). All sequenced samples had RIN values above 7.3 (49).

For dual plant-pathogen transcriptomes *in planta*, samples were harvested 21 days after susceptible cultivar Bonny Best tomatoes were inoculated with ~2000 CFU of each bacterial strain through the cut petiole. 72 h after inoculation approximately 0.1 g stem tissue was collected from the site of inoculation, immediately frozen in liquid nitrogen and stored at −80°C. Another 0.1 g of tissue was collected from directly below the inoculation site, ground in bead beater tubes using a PowerLyzer (Qiagen, Hilden, Germany) for two cycles of 2200 rpm for 90 s with a 4 min rest between cycles. This material was then dilution plated to measure bacterial colonization. Total RNA was then extracted from stem samples colonized with between 10^8^ and 10^9^ CFU/g of tissue using a hot-phenol chloroform method (19). Between 4 and 5 individual plants were pooled per biological replicate. Nucleic acid sample quality was checked using a nanodrop, Agilent bioanalyzer, and qRT-PCR primers Actin_F/R (49). All samples had RIN values above 7.2.

All RNA samples were sent to Novogene (Beijing, China) for cDNA library construction, sequencing, and analysis.

#### Differential expression analysis

Differential expression analysis (for DESeq with biological replicates) was performed using the DESeq R package (1.18.0) (50). DESeq provided statistical routines for determining differential expression in digital gene expression data using a model based on the negative binomial distribution. The Resulting P-values were adjusted using the Benjamini-Hochberg approach for controlling the false discovery rate. Genes with an adjusted P-value of <0.05 found by DESeq were assigned as differentially expressed.

#### R methods

Transcriptional groups of interest were manually selected from GO biological process and cellular function groups. Genes possessing GO annotations referring to multiple transcriptional groups were assigned with priority as follows: Iron, Sulfur, Nitrogen, Oxidative Stress, Cellular Damage, Regulators. Visualization of differential expression using RPMK and log 2-fold change was done in R (version 4.1.0) using the base and graphics packages.

### qRT-PCR Gene Expression

*Rs* cells were grown in 15 ml conical tubes in BD Gaspak anaerobic jars (BD, Franklin Lakes, NJ) for 15 h then 1 mM or 0 M SNP was added and grown for 3 h under hypoxic denitrifying conditions as described above. Total RNA was extracted using a hot phenol chloroform method as described (19). DNA was removed with DNAfree DNase (Invitrogen, Life Technologies, Calrsbad, CA) and cDNA and no-RT controls were synthesized from 200 ng to 1 ug RNA using the SuperScript VILO cDNA synthesis kit (Life Technologies, Carlsbad, CA). The qRT-PCR reactions were run in triplicate with 5 ng cDNA and Power Up SYBR Green Master Mix (Applied Biosystems, Foster City, CA) in a 10 uL volume using an ABI 7300 Real Time PCR System (Applied Biosystems, Foster City, CA). Relative gene expression was calculated using the 2^-ΔΔCT^ method, normalizing to the consistently expressed *rplM* gene (51).

All primer sets amplified fragments between 100-200 bp and had 90-110% efficiency and are listed in Supporting Information Table S5.

### Oxidative Stress Assay

Denitrifying *R. solanacearum* cells were grown in VDM + 30mM NO_3_^-^ in 96-well microtiter plates in anaerobic pouches (BD, Franklin Lakes, NJ) in a 28°C shaking incubator at 225 rpm. After 16 h cells were treated with 0 M or 100 uM Spermine-NONOate and 0 M or 500 uM H_2_O_2_ and returned to pouches with fresh anaerobic sachets for 3 h. After this second incubation, bacterial survival was measured as cell density in a microplate reader (BioTek, Winooski, VT, USA) using absorbance at 600nm (ABS_600_).

### Quantification of intracellular aconitase activity

*Rs* strains were grown overnight in 5 mL VDM at 28°C, 0% O_2_ and cultures were standardized turbidometrically. About 10^10^ CFU were pelleted and resuspended in water with 20 mg/mL lysozyme (Sigma-Aldrich) to a 5 mL volume, then incubated on ice for 45 minutes. Cell suspensions on ice were then sonicated with a needle sonicator at 40% amplification for ten 30 s pulse cycles with 10 s between cycles. The resulting lysates were then used in the aconitase assay (Sigma-Aldrich) in a 96 well plate format according to the kit instructions. Samples were measured at 450 nm in a microplate reader (BioTek, Winooski, VT, USA) and analyzed to determine units of activity per cell according assay protocol.

## Results

### *norA, norB*, and *hmpX* are upregulated in denitrifying cultures and by exogenous NO

The proteins encoded by *norA, norB*, and *hmpX* in *Rs* strain GMI1000 are conserved across diverse bacteria, including environmental isolates and plant and animal pathogens (Table S1A). Further, all three were encoded in genomes of the several hundred sequenced strains in the *R. solanacearum* species complex. Previous functional analyses demonstrated that *Rs norB* encodes an NO reductase and *hmpX* encodes an oxidoreductase (6). We identified locus Rsp0958 as *norA* because its product resembles known single-heme domain proteins that reduce NO and H_2_O_2_ oxidative stress by replacing damaged di-iron centers in Fe-S cluster proteins (36, 40). It is most similar to NorA from *C. necator* (73% AA identity), and to YTFE from *Salmonella enterica* and DnrN from *N. gonorrhoeae* (~50% AA identity), which have been implicated in oxidative stress mitigation (Table S1A). *Rs* NorA, NorB, and HmpX each contain the highly conserved heme or globin metal cofactor binding domains necessary to reduce NO toxicity (Fig. S1). These genomic analyses suggested *Rs* NorA, NorB, and HmpX could all contribute to mitigating oxidative damage.

A previous transcriptomic analysis found that when *Rs* grows in the oxidatively stressful plant host environment, the pathogen upregulates *norA, norB*, and *hmpX* by 75, 51, and 43-fold, respectively, relative to their expression in rich media (19, 31, 45). Indeed, these were among the genes most differentially expressed *in planta* (Table S1B). *norA, norB*, and *hmpX* were also highly expressed in denitrifying *Rs* cells cultured at 0.1% O_2_, a condition that produces an oxidative environment. Treating denitrifying cultures with exogenous NO further increased expression of *norA* (5-fold, *P*= 0.0211, one sample t-test), *norB* (9-fold, *P*=0.0552), and *hmpX* (8-fold, *P*= 0.0255) (Fig. 2). This significant up-regulation of *norA, norB*, and *hmpX* in the oxidative plant environment, and in response to exogenous NO is consistent with the hypothesis that these genes are important for NO metabolism.

**Figure 2.**
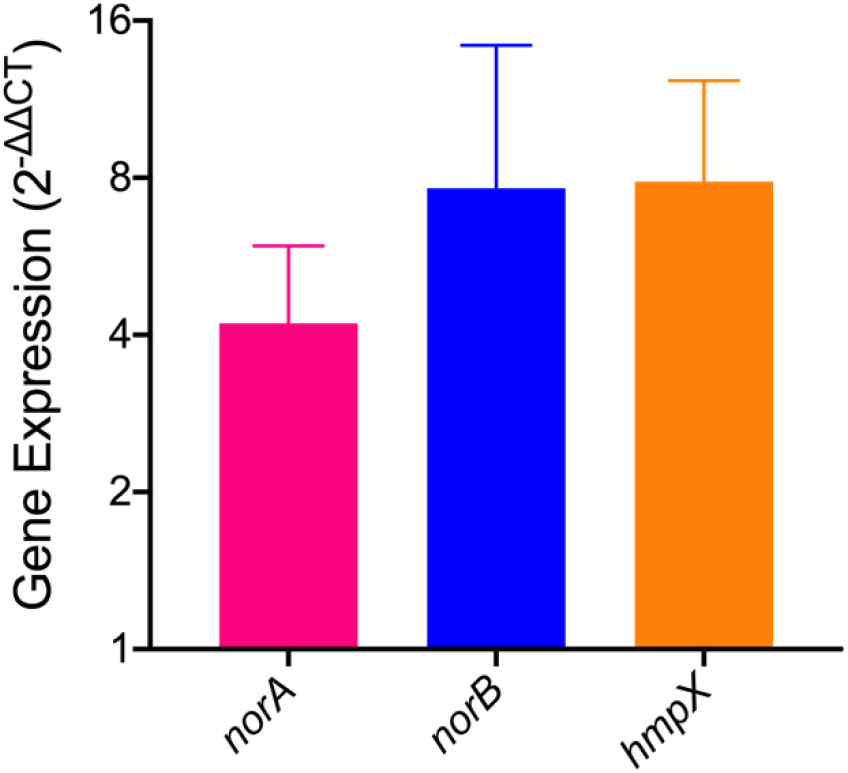
Exogenous NO induces expression of *norA, norB*, and *hmpX*. Relative gene expression, of wild-type *R. solanacearum* GMI1000 as determined by qRT-PCR. RNA was extracted from *R. solanacearum* cells cultured for 16 h under denitrifying conditions (VDM + 30 mM NO_3_^-^ with 0.1% O_2_), then treated with 1 mM NO donor sodium nitroprusside for 3 h in denitrifying conditions. Gene expression is relative to untreated *R. solanacearum* cells. Wild-type gene expression for *norA, norB*, and *hmpX* was normalized to *rplM*. Data are mean +/- SEM (*norA*, P = 0.0211; *norB*, P = 0.0552; *hmpX*, P = 0.0255, one sample t-test). Data are mean of 4 biological experiments, each containing 3 technical replicates. Fold change was calculated using the 2^-ΔΔCT^ method.

To explore whether these three genes are all under the control of the same NO-responsive regulator, we used RegPrecise to find predicted binding sites (52). Binding sites for the NO-responsive Rrf2 family regulator NsrR were present 5’ of *norB* and *hmpX*, but not *norA*. Upstream of *norA* we found a binding site for NorR, the predicted NO-inducible sigma-54 dependent Fnr family regulator. This suggested these genes are under different regulons. In *Rs*, NsrR is predicted to have nine genes in its regulon, but NorR is predicted to regulate only the *norAR* operon (52, 53). However, in other bacteria such as the closely related *C. necator* NorR regulates both *norA* and *norB* (53–56). To confirm the bioinformatic prediction that *Rs* NorR exclusively regulates *norA*, we measured expression of *norA, norB*, and *hmpX* in a *ΔnorR* deletion mutant. Indeed, when *Rs ΔnorR* grew under denitrifying conditions, *norA* expression was reduced 15-fold relative to the wild-type parent strain, while expression of *norB* and *hmpX* did not change (Fig. S2A). This indicates that *norB* and *hmpX* are not regulated by NorR, and that the *Rs* response to NO is complex and involves at least two distinct regulatory mechanisms (Fig S2B). This finding prompted us to investigate the functional interplay of NorA, NorB, and HmpX.

### Δ*norA*, Δ*norB*, and Δ*hmpX* mutants upregulate iron and sulfur metabolism in denitrifying conditions

Oxidative molecules like NO cause nitrosative stress that damages cellular components including Fe-S proteins, lipids, and DNA, leading to the SOS response (39, 57, 58). We hypothesized that cells lacking putative stress mitigation genes *norA, norB*, or *hmpX* would suffer nitrosative damage that would be reflected in altered expression of genes encoding iron, sulfur, and repair pathways. We tested this hypothesis by profiling the transcriptomes of wild-type and Δ*norA*, Δ*norB*, and Δ*hmpX* strains after 16 h growth in denitrifying conditions, a timepoint when NO_3_^-^ respiration generates NO and nitrosative stress. All three mutations substantially affected the *Rs* transcriptional profile. Relative to wild-type *Rs*, the Δ*norA* and Δ*hmpX* mutants had 187 and 281 differentially expressed genes (DEGs), respectively. A surprising 2/3 of the genome, or 4105 of 6200 ORFs, were differentially expressed in the Δ*norB* mutant (Fig. 3).

**Figure 3.**
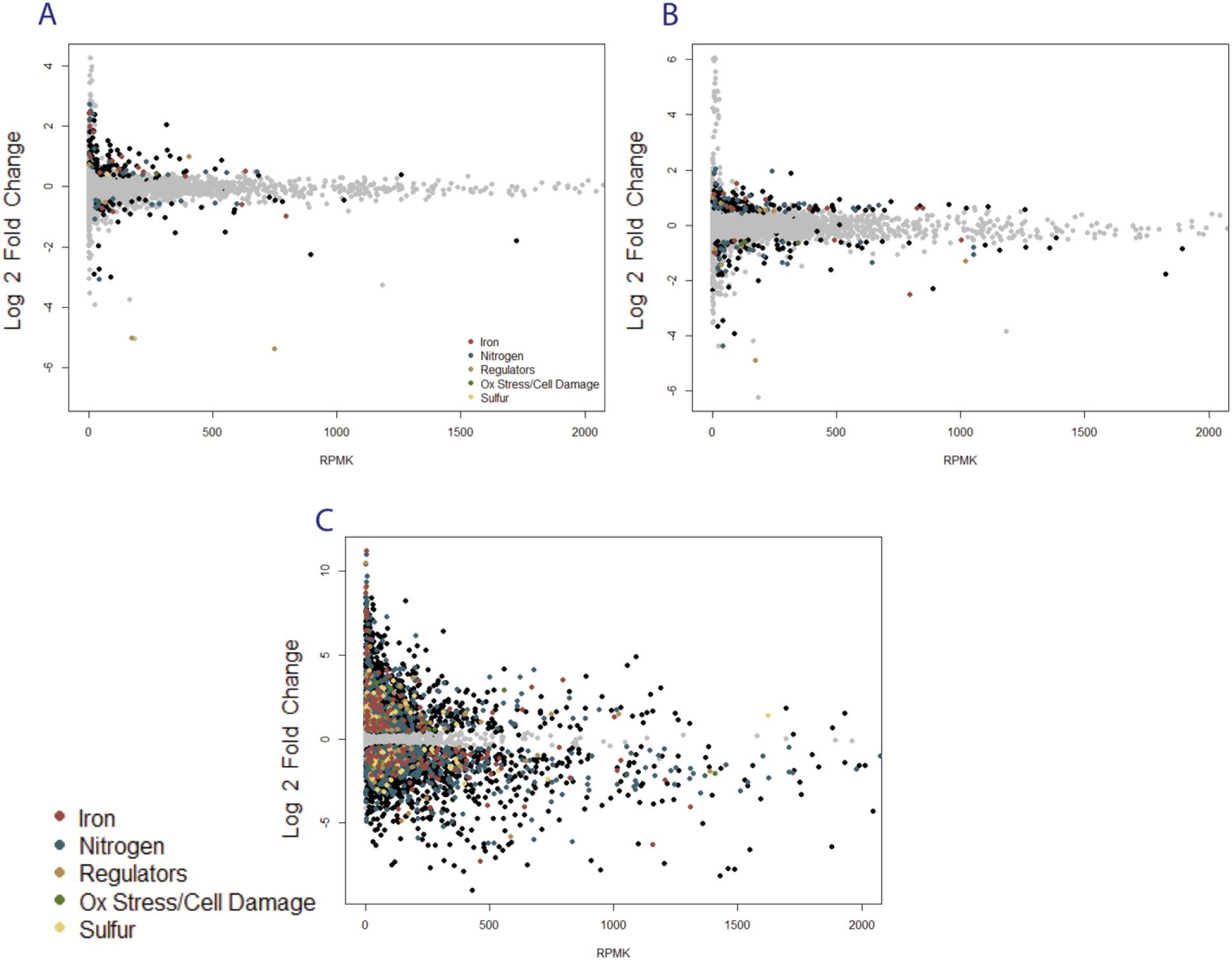
Transcriptomic profiles of *R. solanacearum* Δ*norA*, Δ*norB*, and Δ*hmpX* mutants relative to wild-type. Plots showing gene expression in Δ*norA* (A); Δ*norB* (B), and Δ*hmpX* (C). RNA was extracted and sequenced from bacteria after 16 h growth in denitrifying conditions (VDM + 30 mM NO_3_^-^ with 0.1% O_2_). Log2-fold change in expression is plotted against reads per million per kilobase (RPMK) to show change in regulation vs. transcript abundance, relative to gene expression in wild-type *R. solanacearum* strain GMI1000. Differentially expressed genes (defined as P < 0.05,) are shown in black, except for genes with GO terms related to iron metabolism (red), nitrogen metabolism (blue), regulators (brown), oxidative stress and cellular damage (green), sulfur metabolism (yellow). Genes not differentially expressed are shown in grey. Of the 6108 open reading frames in the *R. solanacearum* genome, Δ*norA* had 187 DEGs, Δ*hmpX* had 281 DEGs, and Δ*norB* had 4112 DEGs compared to wild-type cells.

Many of the 187 DEGs in the Δ*norA* mutant were upregulated and predicted to be involved in stress tolerance, iron acquisition, and inorganic nitrogen metabolism (Fig. 3A). Among the most upregulated DEGs were the iron homeostasis regulator *fur2;* Rsp0415 encoding the putative iron-stress response sigma-factor RpoE; and Rsp0421 putatively encoding RhbC, a component of siderophore synthesis. Among the most abundantly expressed DEGs were: *narG* and *narH* encoding subunits of a nitrate reductase, and Rsc0754 encoding putative peroxidase AhpC. In *ΔnorA, hmpX* was slightly downregulated 1.96-fold (*P* = 3.21E-5) and *norB* expression was not significantly different from wild-type, although it was already in the wild type strain’s top 10 most abundantly expressed genes (Table S2). Overall, this transcriptomic profile suggests that loss of the predicted RIC protein NorA causes increased oxidative stress that affects iron metabolism, but that the *ΔnorA* mutant mitigates this by upregulating genes for a wide range of protective mechanisms.

In the Δ*hmpX* mutant, about half of the 281 DEGs were upregulated and were related to inorganic nitrogen or sulfur metabolism (Fig. 3B). Among the most highly upregulated genes were *nsrR*, encoding a nitrate sensitive repressor; *hsdM* (Rsc3396) and *hsdR* (Rsc3384), encoding a putative type I restriction modification system; and *sbp*, encoding a sulfate binding protein involved in cysteine synthesis. Although *norB* was slightly downregulated in *ΔhmpX* (1.61-fold, *P* = 1.29E-5) and *norA* expression was not significantly different from wild type, both genes remained in the top 20 most abundantly expressed genes, and *norB* was the single most abundant gene transcript expressed by Δ*hmpX* in denitrifying conditions (Table S1B). This profile suggests that Δ*hmpX* is still metabolizing NO and may pivot its metabolic strategies to acquire more sulfur to address damage to iron, sulfur, or Fe-S cluster proteins.

Loss of the NO reductase NorB had the most dramatic transcriptional effect. Genes involved in iron metabolism, sulfur metabolism, or cellular repair were most highly upregulated (Fig. 3C). The top three most upregulated genes, all encoding iron acquisition proteins, were upregulated over 1000-fold (*P* < 3.34E-67). Even the regulator *fur2* was upregulated 854-fold (*P* = 6.26E-89). The *ΔnorA* and *ΔhmpX* transcriptomes showed similar trends but with a smaller magnitude than in *ΔnorB* (Fig. 3C). In addition, *ΔnorB* significantly upregulated *norA* and *hmpX* by 2.43-fold (*P* = 1.38E-9) and 11.67-fold (*P* = 1.9E-58), respectively (Table S1B).

The global up regulation of iron homeostasis regulators like *fur2* in Δ*norA* and Δ*norB* mutants indicated damage to Fe-S cluster proteins, but Δ*hmpX* and Δ*norB* also upregulated error-prone DNA polymerase *dnaE2* 1.62-fold (*P*=0.029) and 118.05-fold (*P*=2.18E-32) respectively, suggesting that cells lacking *hmpX* or *norB* also experience oxidative damage to DNA.

More broadly, mutants lacking either *norA, norB*, or *hmpX* shared 43 common DEGs, 21 of which have known homologs or domains with predicted function (Fig. 4). All three mutants differentially expressed bacterioferritin-encoding *bfd* and seven genes related to sulfur metabolism. Further, all three mutants upregulated *paaE*, which is predicted to encode degradation of phenylacetic acid (PAA) or a plant auxin growth hormone, which could interact with plant hosts. Interestingly, the most downregulated genes for all three mutants were in the Rsp1617-1623 operon (about 10-30-fold, *P* < 0.021949), which is predicted to be involved in cell attachment. Together these shared DEGs suggest that all three mutants suffer enough RNS to cause detectable cellular damage.

**Figure 4.**
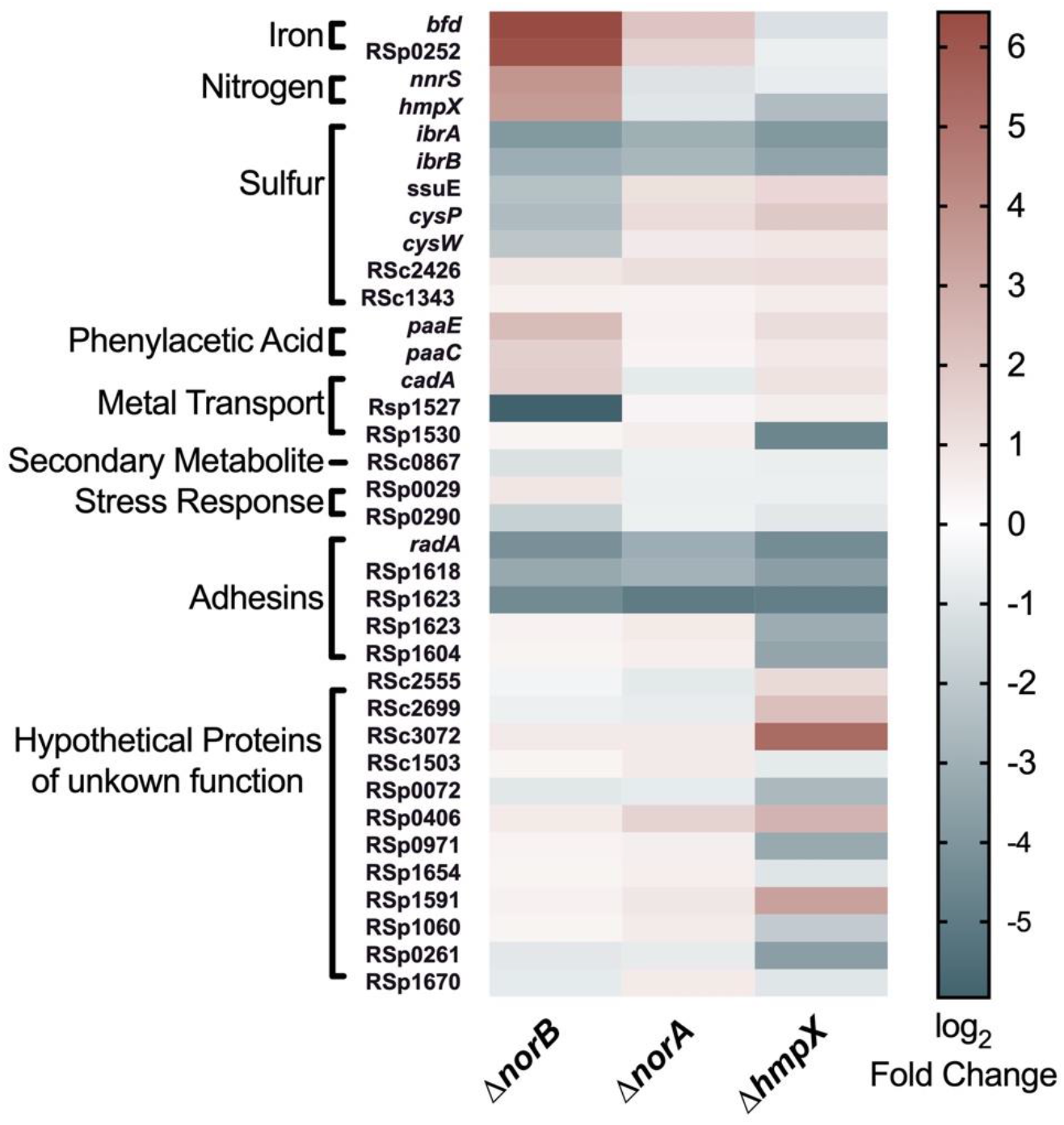
*R. solanacearum* mutants lacking NorA, NorB, and HmpX differentially expressed some of the same genes. RNA was extracted and sequenced from bacteria after 16 h growth in denitrifying conditions (VDM + 30 mM NO_3_^-^ with 0.1% O_2_). Differentially expressed genes (DEGs, relative to wild-type strain GMI1000) from all three mutants were compared, and shared DEGs were joined with SQL. Known or putative function was used to sort genes into categories labeled on the left. Log2 fold change is represented as a heat map showing expression of selected shared DEGs relative to expression levels to wild-type cells. Red indicates upregulated genes, white indicates genes not significantly different from wild-type, and blue indicates downregulated genes, as shown in the scale bar at right.

### A mutant lacking *norB* accumulates NO in culture, and has severely reduced virulence *in planta*

Transcriptomic analysis suggested that *norA, norB*, and *hmpX* are important for mitigating RNS stress that *Rs* experiences during denitrifying respiration in culture and in the low-oxygen plant host xylem (6, 19). We directly tested this hypothesis by assessing in culture and *in planta* behaviors of *Rs* deletion mutants lacking *norA, norB*, or *hmpX* (Fig. 5).

**Figure 5.**
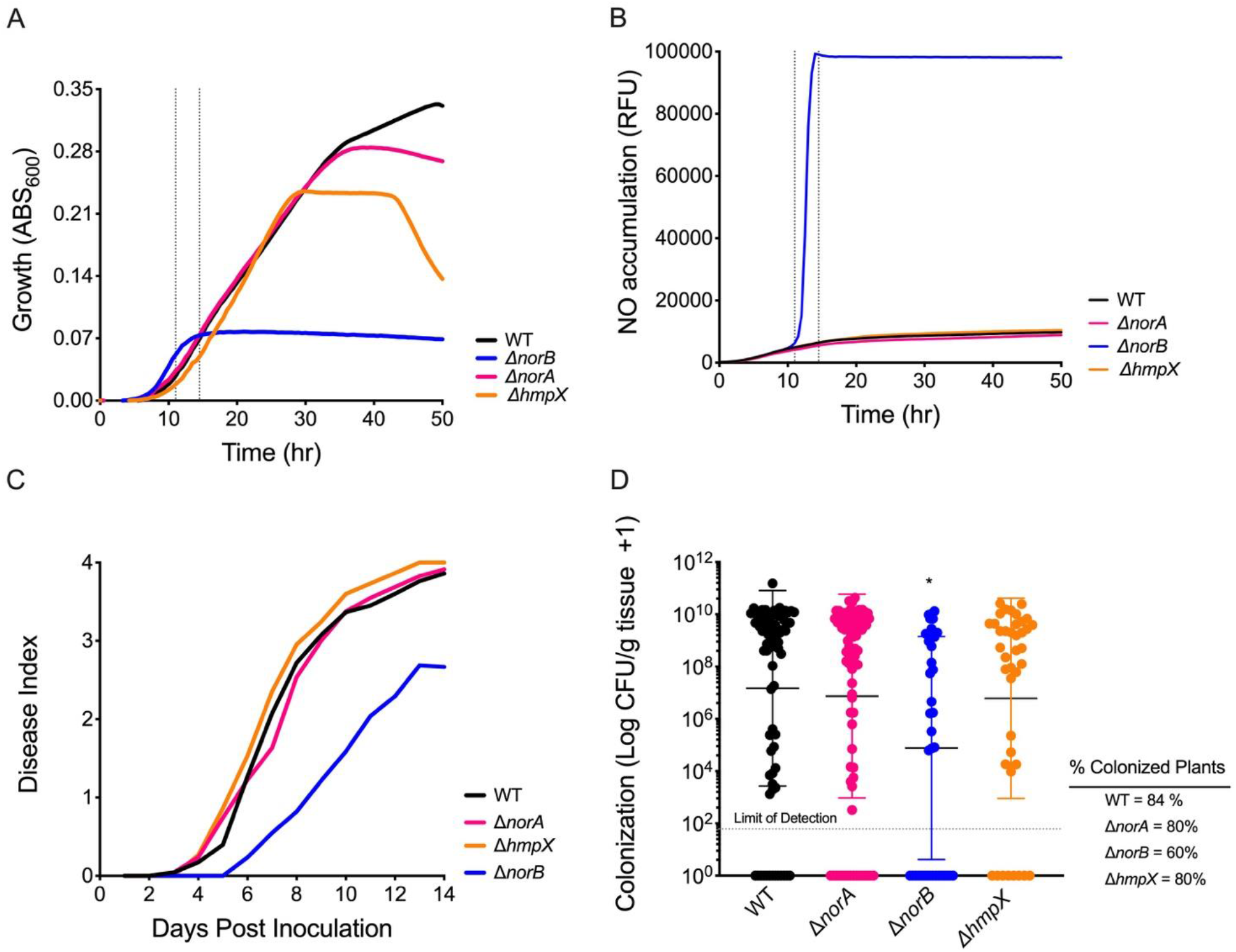
Behavior of *R. solanacearum* Δ*norA*, Δ*norB*, and Δ*hmpX* mutants in denitrifying culture conditions and *in planta*. **A.** Growth of wild type and mutant *R. solanacearum* cells in denitrifying conditions (VDM + 30 mM NO_3_^-^ with 0.1% O_2_) in shaking 96 well plates, shown as Abs _600_. Data are mean+/- SEM Data and are mean of 4 biological experiments, each with 3 technical replicates. Gray bars represent time of toxic NO accumulation ~12 h. **B.** Accumulation of nitric oxide (NO) over time in the cultures in panel A, measured as relative fluorescence units using the NO-specific fluorescent indicator DAF-FM-DA. Excitation and emission measured at 495/515 nm. Data are mean of 4 biological experiments, each with 3 technical replicates. Gray bars represent time of toxic NO accumulation ~12 h.**C.** Bacterial wilt disease progress on 16-day-old wilt-susceptible ‘Bonny Best’ tomato plants following naturalistic soil-soak inoculation with 1×10^8^ CFU wild-type or mutant *R. solanacearum* cells. Plants were assessed for wilt symptoms on a scale of 0-4 over 14 days. Data shown represent the mean disease index of 45-93 plants in 3-6 biological replicates. Virulence of the Δ*norB* mutant was lower than that of the three other strains (*P* = 0.0018, Repeated Measures ANOVA). **D.** *R. solanacearum* population sizes in tomato mid-stems 4 days after 2×10^6^ CFU of *R. solanacearum* were applied to the cut petiole of the first true leaf. Bacterial populations were quantified by grinding and serially dilution

In aerobic culture, when *Rs* does not denitrify, the Δ*norA*, Δ*norB*, and Δ*hmpX* strains grew as well as parent strain GMI1000 (Figure S3). None of the three mutants grew as well as wild type in hypoxic denitrifying culture, although their growth was affected to differing degrees (Fig. 5A). For the first 24 h, Δ*norA* and Δ*hmpX* grew like wild type, but their growth plateaued at ~36 and ~28 h, respectively, while wild type did not enter stationary phase until ~48 h. Growth of the Δ*norB* mutant under denitrifying conditions plateaued much earlier at ~12 h, while wild type was still in early log phase growth. The limited growth of the *ΔnorB* mutant was consistent with development of toxic conditions that interfered with bacterial growth.

To directly test whether these three mutants accumulate NO, we used the NO-specific fluorescent probe DAF-FM-DA to measure NO accumulation over time in denitrifying cultures (Fig. 5B). The ~12 h growth plateau of the Δ*norB* mutant correlated exactly with a rapid accumulation of NO in the culture, which contained at least 10 times more NO than wild-type cultures. Wild type, Δ*norA*, and Δ*hmpX* cells did not accumulate detectable amounts of NO, likely because NorB reduces NO almost as fast as it is produced in all three strains.

We previously determined that Δ*norB* has a virulence defect, and that neither *ΔnorB* nor Δ*hmpX* colonize tomato plants as well as wild-type following a naturalistic soil-soak inoculation (6). To see if loss of *norA* also affected these behaviors, we inoculated tomato plants with either Δ*norA*, Δ*norB*, or Δ*hmpX*. The *ΔnorB* mutant caused significantly reduced bacterial wilt symptoms in the soil-soak assay (Fig 5B). By 72 h after tomato stems were directly inoculated through a cut leaf petiole the population size of *ΔnorB* in tomato mid-stems was around two orders of magnitude smaller than that of wild type (Fig. 5C). In contrast, neither the Δ*norA* nor the *ΔhmpX* mutant differed significantly from wild type with respect to bacterial wilt virulence or stem colonization after petiole inoculation. Results of these *in planta* experiments are consistent with the finding that the Δ*norB* mutant accumulates toxic levels of NO that severely impair its growth in denitrifying culture. In contrast, the Δ*norA* and Δ*hmpX* mutants functioned much like wild type in both conditions. The *in planta* defects of *ΔnorB* are likely explained by the mutant’s inability to detoxify the NO generated by denitrifying respiration during plant pathogenesis. The defects further suggest that without either NorA or HmpX, *Rs* can overcome oxidative stress produced by bacterial denitrification and the plant host, likely by changing the transcription of iron and sulfur metabolism genes. However, despite massive transcriptomic changes *Rs* cannot compensate for loss of the NorB nitric oxide reductase as evidenced by the mutant’s loss of virulence, plant colonization defects, and reduced fitness in culture.

### NorA, NorB, and HmpX function together in denitrifying culture

Detoxification of reactive radical species like NO is critically important for fitness of denitrifying bacteria (59–61). Although Δ*norA* and Δ*hmpX* single mutants had wild-type virulence and were only modestly reduced in late stage denitrifying growth compared to wild-type *Rs*, their transcriptional signatures indicated they did suffer RNS stress early in denitrifying cultures. Additionally, during denitrification the Δ*norB* mutant strongly upregulated expression of *norA* and *hmpX*. We wondered how *Rs* would behave in the absence of two or more components of its RNS mitigation system.

We therefore created double deletion mutants lacking multiple genes; *norA* and *norB (ΔnorAB); norA* and *hmpX (ΔnorAX); hmpX* and *norB (ΔnorBX)*. Persistent efforts to use the same methods to create a *ΔnorA/norB/hmpX* triple mutant were unsuccessful, suggesting that loss of all three proteins is lethal to *Rs*. After 16 h of growth under denitrifying conditions (corresponding to the time RNA was harvested for transcriptional analysis), the Δ*norAX* double mutant grew as well as wild type. However, both double mutants lacking *norB* grew to lower endpoints (yield) than WT, Δ*norA*, or Δ*hmpX (P* < 0.0078, ANOVA), although the growth of Δ*norAB* and Δ*norBX* was not significantly different from that of the Δ*norB* single mutant (Fig. 6A). After 36 h under denitrifying conditions, all single and double mutants had significantly lower Abs_600_ than wild type. Further, single and double mutants lacking NorB were dramatically reduced in growth at 36 h (Fig. 6B). The Δ*norAB* and Δ*norBX* double mutants grew only around 10% as much as wild-type, Δ*norA*, or Δ*hmpX* cells (*P* < 0.001, ANOVA). Additionally, these double mutants also reached a 35% lower Abs_600_ reading than the Δ*norB* single mutant (*P* < 0.001, ANOVA). These cumulative growth differences show that the NorB nitric oxide reductase plays an irreplaceable role in mitigating NO stress both early and late in denitrifying growth in culture. However, the RIC protein NorA and oxidoreductase HmpX also protect *Rs* when NO accumulates, especially during later stages of denitrifying metabolism.

**Figure 6.**
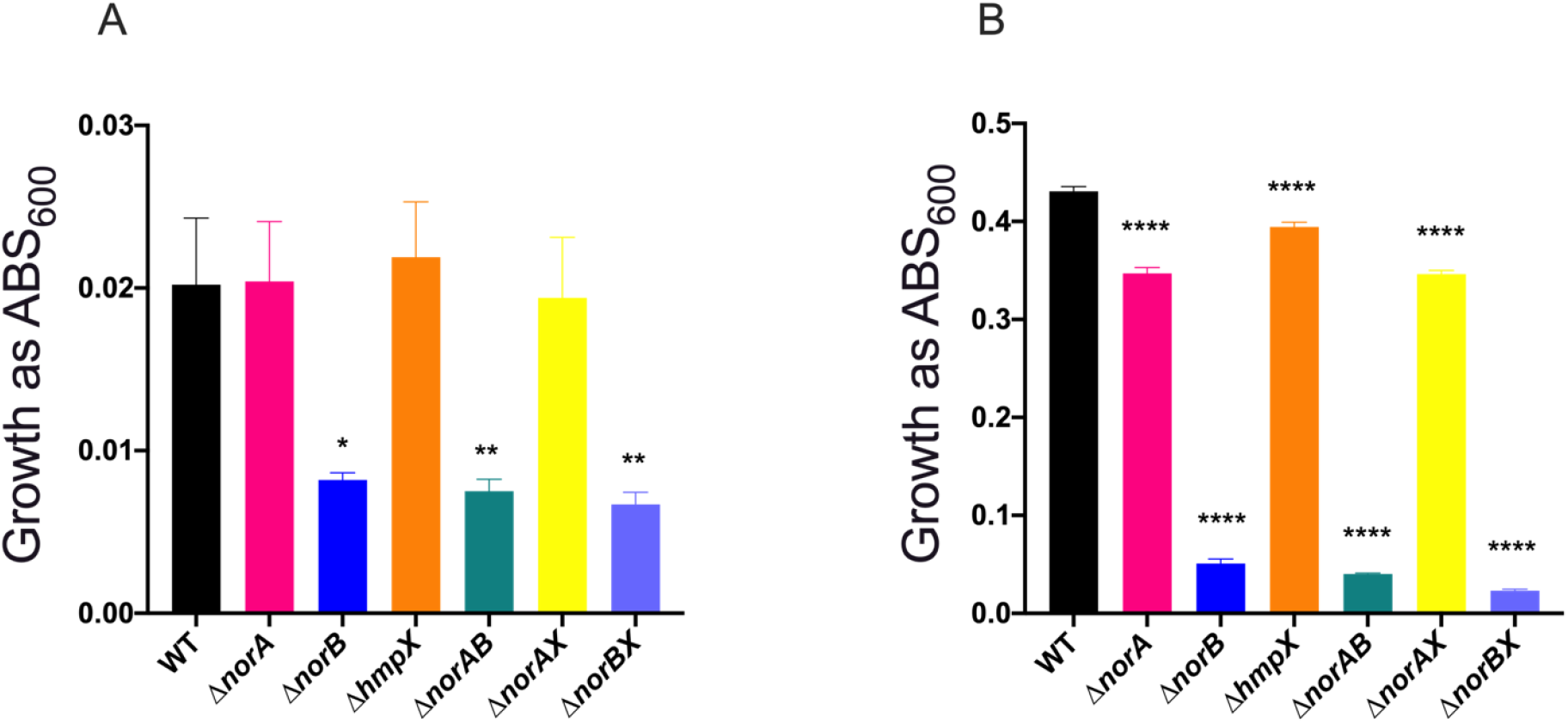
The NorB nitric oxide reductase is important for *R. solanacearum* growth in denitrifying conditions. Growth of wild type, Δ*norA*, Δ*norB*, Δ*hmpX*, and double mutants in VDM + 30 mM NO_3_^-^ with 0.1% O_2_ shaking in 96 well plates, measured spectrophotometrically as absorbance at 600 nm, after **A.** 16 h culture or **B.** 36 h culture. Data shown reflect the mean +/- SEM of 4 biological replicate experiments, each containing 3 technical replicates. For both 16 h and 36 h, asterisks indicate difference from growth of the wild-type strain: * P < 0.05, **P < 0.008, **** P < 0.001 (one-way ANOVA).

### *norA, norB*, and *hmpX* contribute to cellular protection from oxidative stress

Denitrifying metabolism damages iron-sulfur (Fe-S) cluster proteins like the TCA cycle enzymes fumarase and aconitase by binding to iron and changing the oxidative state of the bound catalytic center (36, 62). *Rs* mutants lacking *norA, norB*, or *hmpX* altered expression of many genes involved in iron and sulfur metabolism, which suggested these mutants experienced damage to Fe-S proteins and would be more susceptible to oxidative stress.

To test the hypothesis that *Rs* lacking NorA, NorB, or HmpX are more susceptible to oxidative stress, we treated denitrifying cultures with exogenous NO or H_2_O_2_ at 16 and 36 h, then measured their growth recovery (Fig. S4). At 16 h, all tested strains recovered similarly from exposure to the NO donor spermine-NONOate (Fig S4A). At 36h, the Δ*norB* mutant actually recovered from NO treatment better than all other strains *(P*<0.0001, ANOVA). (Fig. S4B). Similarly, the Δ*norAB, ΔnorAX*, and *ΔnorBX* double mutants were more tolerant of H_2_O_2_ than wild type at 16 h (Fig. S4C), although their recoveries did not differ at 36 h (Fig. S4D). We concluded that single or double mutants lacking *norA, norB*, or *hmpX* were not more susceptible to the levels of exogenous oxidative stress tested under these conditions.

As a measure of Fe-S cluster damage, we quantified aconitase activity in various *Rs* strains growing in denitrifying conditions, normalizing enzyme activity to cell density to account for differences in growth between strains. After 16 h of culture, wild-type and all mutant cells contained similar aconitase levels (data not shown). However, by 36 h, all strains lacking *norB* had reduced aconitase activity compared to wild-type cells (Fig. 7). While the wild-type strain contained an average of 0.58 milliunits/mL, Δ*norB*, Δ*norAB*, Δ*norBX* produced 0.39, 0.33, and 0.27milliunits/mL of active aconitase respectively (*P* = 0.0360, 0.0057, and 0.008 respectively, ANOVA). Aconitase activity in Δ*norAB* and Δ*norBX* double mutants trended lower than that in the *ΔnorB* single mutant, although they were not significantly different. At 0.42 milliunits/mL, aconitase activity in the Δ*norAX* mutant similarly trended down but was not significantly different from wild-type. Together with the transcriptional profiles suggesting that *ΔnorA* and *ΔhmpX* experience iron and sulfur stress, these trends indicate that NorA and HmpX help protect Fe-S proteins, including aconitase. However, NorB is the major source of *Rs* cellular protection in denitrifying conditions, as evidenced by both transcriptional and direct enzyme analyses.

**Figure 7.**
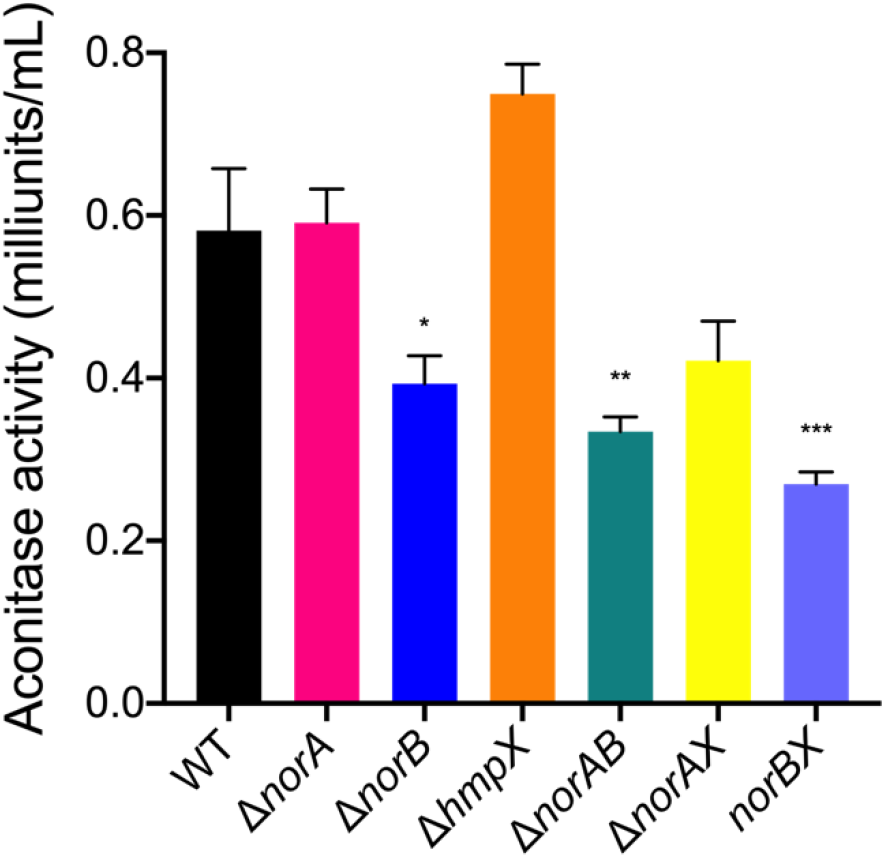
*R. solanacearum* needs the NorB nitric oxide reductase to prevent damage to the iron-sulfur protein aconitase. *R. solanacearum* wild type, Δ*norA*, Δ*norB*, Δ*hmpX*, and double mutants were cultured for 36 h in VDM + 30 mM NO_3_^-^ with 0.1% O_2_. For each strain, activity of the abundant iron-sulfur protein aconitase was measured using the Aconitase Activity Assay Kit (Sigma-Aldrich) according to the manufacturer’s instructions. Data shown reflect the mean+/- SEM of 3 biological replicates. Asterisks indicate difference from growth of the wild-type strain: * P = 0.0360, **P = 0.0057, *** P = 0.0008 (ANOVA).

### Bacterially-produced NO affects plant host transcriptional responses

Having shown that oxidative stress is toxic to *Rs* cells both *in planta* and in culture, we investigated ways ROS could affect bacterial-plant interactions. NO is a free radical signaling molecule that affects every stage of the plant life cycle (27, 63, 64). In particular, NO interacts with plant hormones to change signaling pathways during plant growth and biotic interactions (27, 63, 65). We hypothesized that increasing the amount of NO produced by the pathogen would alter plant perception of *Rs* during infection. We tested this by comparing the transcriptomes of tomato plants infected with either wild-type *Rs* or the NO-accumulating Δ*norB* mutant to the transcriptome of healthy plants. As expected, in response to infection by either wild-type *Rs* or *ΔnorB*, tomato plants significantly changed gene expression patterns, including pathways in the KEGG and GO categories of general cellular metabolism and processes involved in plant-pathogen interactions. (Fig. 8, Fig S6, Table S4). Differentially expressed genes (DEGs) fell into 39 KEGG categories in plants infected with wild type *Rs* and 42 categories for Δ*norB*-infected plants, with 34 KEGG categories shared by plants infected with either strain. Overall, Δ*norB* induced about twice as many DEGs in tomatoes as wild type *Rs* (Fig 8A). Most DEGs in plants infected with either wild-type or Δ*norB* mutant cells changed expression of basic metabolic pathways, biosynthesis of secondary metabolites, plant-pathogen response, and plant hormone signal transduction. Wild-type *Rs* induced more DEGs involved in tomato starch and sucrose metabolism and photosynthesis. While wild-type *Rs* induced plant hormone signal transduction, Δ*norB* mutant cells suppressed plant hormone signal transduction. Wild-type and Δ*norB* mutant uniquely expressed plant DEGs in 5 and 8 KEGG categories respectively (Fig. S6A). Specifically, wild-type cells upregulated host plant nitrogen metabolism and carotenoid biosynthesis, Δ*norB* cells induced biosynthesis of arginine and alkaloids.

**Figure 8.**
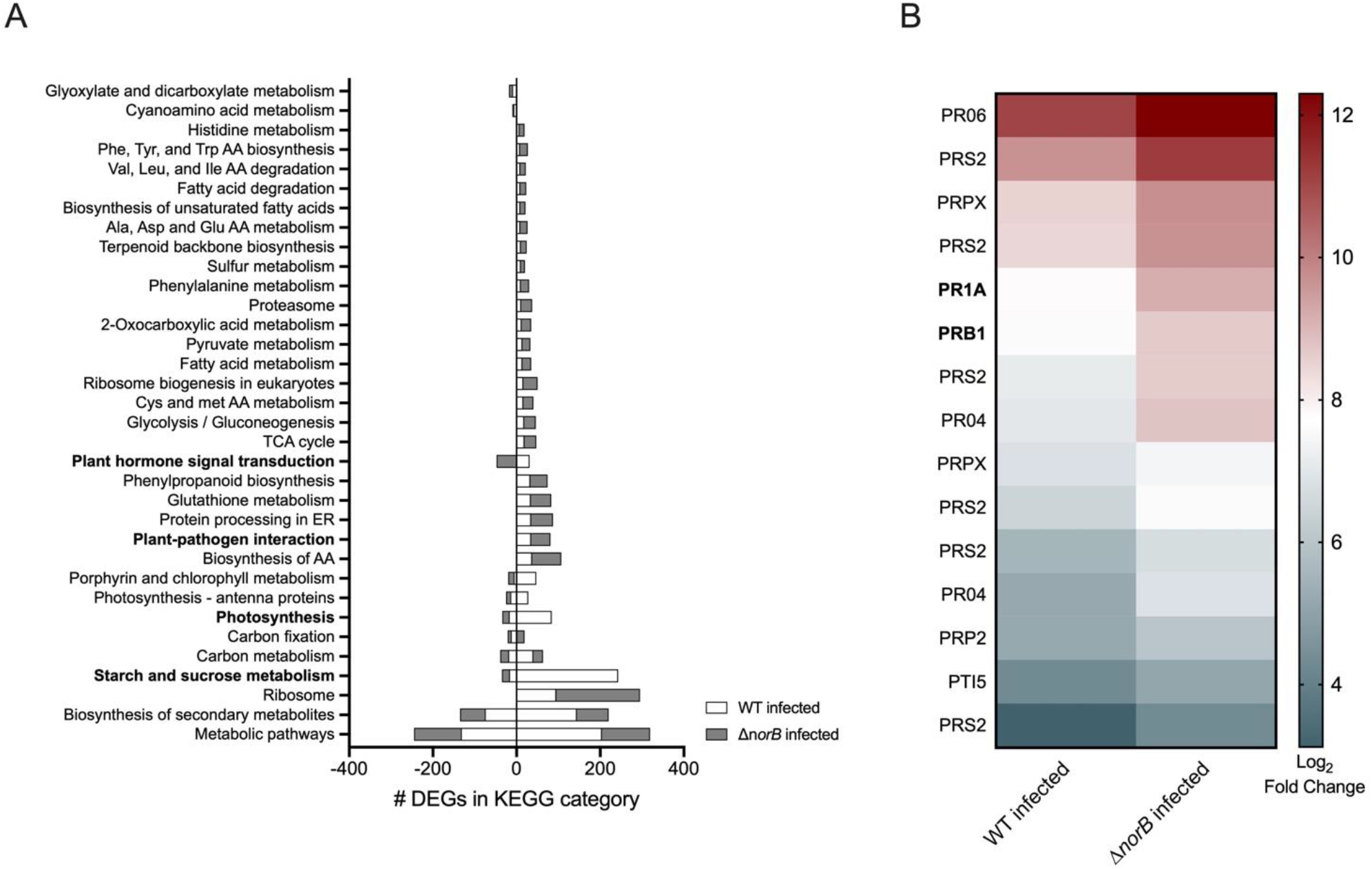
Transcriptomic response of tomato plants infected with either *R. solanacearum* wild-type strain GMI1000 or the Δ*norB* mutant. RNA was harvested and sequenced from stems of ‘Bonny Best’ tomato plants 72 h after they were petiole-inoculated with 2×10^6^ CFU of either wild-type *R. solanacearum*, Δ*norB* or water as a healthy control to determine differential gene expression. **A.** Differentially expressed tomato genes (DEGs) were sorted into 34 KEGG categories shared by both wild-type and Δ*norB* infected plants. Number of DEGs in each KEGG category from wild-type or Δ*norB* infected plants are graphed in stacked columns for comparison. The X axis shows the number of DEGs assorted into the indicated KEGG pathways, as determined with KOBAS software by NovoGene. For details on identification of DEGs, see methods. **B.** Expression levels of 15 pathogenesis-related genes in plants infected with either wild-type or Δ*norB R. solanacearum* cells. Pathogenesis related DEGs were identified by searching the uniprot annotations (https://www.uniprot.org/blast/) of all DEGs for wild-type and Δ*norB* infected plants for any genes annotated with the terms pathogen, pathogenesis, or biotic interaction.

Most strikingly, plants infected with *ΔnorB* differentially upregulated all the pathogen response (PR) genes annotated with the KEGG terms pathogen, biotic, and defense, including the salicylic acid and ethylene pathway defense signaling genes PR1a and PR1b that were previously validated as contributing to tomato resistance to bacterial wilt (Fig 8B). Together, the KEGG and GO-term analyses of tomato DEGs showed that plants had different transcriptional responses to infection by wild-type *Rs* and Δ*norB*. In particular, the tomato host mounted stronger defenses against the NO-overproducing Δ*norB* mutant, possibly because the higher NO levels activate plant defense signaling pathways.

## Discussion

Few bacteria can compete in the low-nutrient, low-oxygen niche of plant xylem vessels, but *R. solanacearum* (*Rs*) thrives in xylem in part by respiring on nitrate. The disadvantage of this metabolic strategy is that it generates potentially toxic levels of highly reactive NO as a byproduct. In addition, *Rs* cells in xylem confront ROS and RNS released by plant defenses (31, 51). Our goal was to determine how this pathogen protects itself from the resulting oxidative stress (6, 19). These mechanisms have been well studied in human pathogens, but little is known about how plant pathogenic bacteria mitigate the damaging effects of oxidative conditions they encounter in their hosts (31, 45).

Many bacteria accomplish this task with nitric oxide reductases (NORs) like NorB, flavorubredoxin oxido-reductases like HmpX, and repair of iron center proteins (RICs) like NorA (36, 66). *Rs* homologs of all three of these proteins were well conserved at the amino acid level, notably at residues that bind cofactors.

The NorA hemerythrin-like domain includes the histidine residues needed to bind the iron cofactor, which are likely responsible for its RIC activity (62). We found that *norA*, but not *norB* or *hmpX*, is regulated by the NO-inducible transcriptional regulator NorR. Transcriptomic analysis of a *ΔnorR* mutant indicated that *norA* is the only enzyme-encoding gene in the NorR regulon; this is noteworthy because NorR typically also regulates *norB* and/or *hmpX* (53, 55). The roles and regulation of RIC/NorA homologs have been studied in a few human pathogens but have not been considered in a plant pathogen (8, 28, 36, 40, 44).

NorB contains a large well conserved heme-copper oxidase domain responsible for NO reductase activity; this domain had homology to many other NOR proteins (12). Single subunit membrane-bound NORs like NorB are typically tied to the electron transport chain and generate ATP (12). However, rapid accumulation of NO in the Δ*norB* mutant made it impossible to distinguish phenotypic effects of energy loss from those of NO toxicity, or to experimentally determine whether NorB contributes to ATP generation in *Rs*.

HmpX, which needs O_2_ for its NO oxidase activity, can also reduce NO in anoxic conditions. The fact that HmpX contains highly conserved residues in both the globin-like NO-binding domain and in the FAD/NAD binding domains needed for full oxidoreductase activity suggests that *Rs* denitrifies or encounters RNS stress in both microaerobic and anoxic conditions (38). Both conditions occur in xylem vessels of *Rs*-infected plants (6). In addition to encountering low oxygen in plant hosts, *Rs* likely experiences low-oxygen denitrifying conditions in soil during its saprophytic life between plant hosts. Many soil-dwelling microbes depend on nitrate respiration and denitrification to thrive in highly variable soil microenvironments (67).

Taken together, the high conservation of these three protective proteins not only in *Rs*, but in other pathogens that do not contain the full denitrification pathway, such as enteric pathogens *E. coli* and *S. enterica*, suggests they are important for pathogen-host interactions, possibly to mitigate oxidative host defenses (68–70).

Transcriptomic analysis of *Rs* during denitrification revealed that NO damage globally changes the bacterium’s gene expression. NorB and HmpX were recently shown to help *Rs* colonize tomato plants, but it was not clear if they contribute to *in planta* fitness because they mitigate nitrosative stress. Wild-type cells treated with NO strongly upregulated *norA, norB*, and *hmpX*, suggesting an important role in nitrosative stress response. Further, *Rs* mutants lacking these three genes had transcriptional signatures consistent with oxidative stress. In denitrifying conditions, all three mutants upregulated iron and sulfur metabolism to varying degrees. However, the Δ*norA*, Δ*norB*, and Δ*hmpX* mutants also had distinct transcriptional profiles and they differentially expressed some shared DEGs at different magnitudes, suggesting redundant functionality by distinct mechanisms and a hierarchical importance where NorB > HmpX > NorA.

The most differentially expressed and most abundant gene transcripts in the *norA, norB, and hmpX* mutants were associated with iron and sulfur metabolism, consistent with damage to Fe-S proteins caused by accumulated nitrosative stress (36, 39, 71). Because NO is both highly reactive and diffusible, it harms many cellular components and can also interact with S-nitrosylated proteins to change transcription in both the bacterium and the plant (72). The catalytic centers of iron and sulfur proteins are especially susceptible to oxidative damage (73). Common bacterial responses to nitrosative stress and Fe-S damage include upregulation of iron sulfur cluster biogenesis genes *isc/nif*, siderophore biosynthesis and secretion, and general bacterial stress response (SOS) systems (73–75). The transcriptomes of denitrifying *Rs* strains were consistent with this pattern. The Δ*norA* mutant upregulated an iron sulfur biogenesis operon including *iscS/R* (Rsc1018-1026), and many iron acquisition genes, including the major ferric uptake regulator FUR2 and putative siderophore biosynthesis and receptor proteins, Rsp0419 and Rsp0416. This suggests NorA normally mitigates oxidative stress by repairing iron centers, so in its absence *Rs* cells become iron limited. The Δ*hmpX* mutant upregulated sulfur metabolism including *ssuB/E* and *sbp* genes, as well as the error prone DNA polymerase *dnaE2*. Upregulation of sulfur and damage response proteins is consistent with upregulation of sulfur metabolism to re-generate or repair damaged bio-available sulfur in Fe-S centers (76, 77). Alternatively, *ΔhmpX* may acquire more sulfur to repair cysteine, which is commonly destroyed by oxidative stress (78). Over 2/3 of the *Rs* GMI1000 genome was differentially expressed in the Δ*norB* mutant, which suffered intense nitrosative stress. As observed for *ΔnorA* and Δ*hmpX*, many of this mutant’s most upregulated and most abundantly expressed genes were involved in iron and sulfur metabolism, but *ΔnorB* also upregulated additional damage response pathways. The Δ*norB* mutant transcriptome carries the signatures of substantial NO damage and an oxidative stress response, consistent with its growth defects in denitrifying culture and *in planta*.

Intriguingly, all three single mutants downregulated a cluster of genes encoding putative collagen-like binding adhesins. These are likely involved in cell-to-cell or cell-to-host attachment. Suppression of adhesion-related proteins suggests the hypothesis that *Rs* cells respond to oxidative stress by detaching from fellow bacteria or xylem vessel surfaces. Stress-induced detachment could help *Rs* cells escape from dense biofilms where toxic levels of NO accumulate, or from host cells releasing oxidative bursts. All three single mutants also upregulated degradation of the auxin phenylacetic acid, a plant growth hormone; auxins help shape tomato defenses against *Rs* (79, 80). By reducing levels of a plant hormone, *Rs* could change plant signaling, and reduce the oxidative defense response. It would be interesting to determine if a *Δpaa* deletion mutant of *Rs* is less successful in plant hosts.

We previously determined that NorB acts in denitrifying conditions such as those found in xylem, but a mutant lacking this enzyme was as virulent as wild type when it was introduced directly into tomato xylem through a cut leaf petiole (6). However, deleting *norB* did significantly lower *Rs* virulence in a more holistic soil soak inoculation assay that forces the pathogen to find, enter, and colonize unwounded plants through the roots. Reduced Δ*norB* mutant virulence following this naturalistic inoculation method suggests that *Rs* depends on NorB during the plant invasion process. At this point *Rs* cells may be more susceptible to oxidative stress produced by other *Rs* cells, competing microbes, or by the plant host. Although the Δ*norA* and Δ*hmpX* mutants had wild-type virulence and plant colonization, our *in vitro* experiments confirmed that NorA, NorB, and HmpX are all required for normal growth under denitrifying conditions. Although Δ*norA* and *ΔhmpX* strains suffered only mild growth defects in denitrifying culture, these two proteins may be important for NO detoxification in the microaerobic soil environments where *Rs* survives between plant hosts. It would be interesting to see if the Δ*norA*, Δ*norB*, and Δ*hmpX* mutants survive as well as wild-type *Rs* in low-oxygen soil microcosms.

Growth of single Δ*norA* and Δ*hmpX* mutants in denitrifying culture plateaued earlier than that of wild type and furthermore these mutants had significant growth defects at 36 h but not 16 h, suggesting these proteins contribute to *Rs* fitness when oxidative stress accumulated. Under these conditions the Δ*norB* mutant quickly accumulated large amounts of NO, and its growth arrest coincided exactly with spiking NO levels in the culture. The toxic effects of NO likely drove the global gene expression changes observed in the Δ*norB* mutant, which was sampled for transcriptomic analysis after 16 h of culture. These data suggest that at this point Δ*norB* cells were so damaged they were simultaneously trying to repair proteins and synthesize them *de novo*. In an apparent attempt to compensate, the Δ*norB* mutant also upregulated expression of *hmpX* and *norA*, as well as genes for many Fe-S enzymes including aconitase. Enzyme activity assays confirmed that *Rs* strains lacking *norB* had reduced aconitase activity, a direct indicator of global cellular damage. In contrast, aconitase activity was not significantly lower than wild type in *norA* or *hmpX* single or double mutants. This suggests that cells depend on NorA and HmpX when NorB can no longer reduce the cellular pool of NO. Measuring growth of Δ*norA* and Δ*hmpX* mutants on older plants that have more developed immune systems, larger xylem vessels, and larger populations of denitrifying bacteria where the pathogen experiences more oxidative stress per cell could reveal if NorA and HmpX make quantitative fitness contributions in late stage disease.

We hypothesized that loss of RNS mitigating proteins would make *Rs* more susceptible to oxidative stress, but on the contrary, all three double mutants trended towards increased ability to recover from treatment with H_2_O_2_. We speculate that because of their defects, these strains were already experiencing enough oxidative stress that they were primed to mitigate the inhibitory effects of H_2_O_2_ more effectively than wild-type (81). This is consistent with our previous observation that *Rs* cells isolated directly from the oxidative plant environment have higher tolerance of oxidative and cold stress than *Rs* cells grown *in vitro* (51). Analyzing the transcriptomes of double mutants could reveal if their unexpectedly high stress tolerance is explained by upregulation of genes involved in iron and sulfur metabolism, the SOS response, and other stress repair mechanisms.

Tomato plants responded differently at the transcriptional level to infection with NO-accumulating Δ*norB* mutant than to infection with wild-type *Rs*. Relative to healthy control plants, *ΔnorB* induced more tomato DEGs than wild type *Rs*. However, plants infected with wild-type *Rs* expressed more starch and sucrose metabolism genes and more genes involved in photosynthesis. This could indicate that during successful infection *Rs* cells manipulate their plant host to increase available nutrients. It is theorized that *Rs* forces plants to load sugar into xylem sap, but the mechanism is still unknown (46). Alternatively, increased defenses triggered by the *ΔnorB* mutant may reduce photosynthesis as part of the well-established growth versus defense tradeoff. It was also interesting that only wild-type *Rs* differentially induced genes in the KEGG category “nitrogen metabolism”. However, arginine biosynthesis was upregulated exclusively in Δ*norB* infected plants. Arginine is thought to be involved in plant nitic oxide synthase (NOS) activity, which oxidizes L-arginine to NO and L-citrulline (82). Increased arginine expression by Δ*norB* infected plants suggests that either the NO accumulated by this mutant is sufficient to change plant signaling and induce NOS, or that accumulated NO is causing a damage response.

Plants can recognize damage-associated molecular patterns (DAMPs, such as cell wall fragments and extracellular non-self-DNA) and pathogen-associated molecular patterns (PAMPs, like flagellar proteins and peptidoglycan) (83–85). In response to DAMPs and PAMPs, both plants and animals produce a defensive burst of ROS and RNS like H_2_O_2_ and NO (86). As discussed above, its strong oxidative properties make NO a potent antimicrobial compound. However, NO is also a key actor in plant defense signaling pathways. Notably, all tomato genes annotated with the terms pathogen, biotic, and defense were expressed at higher levels in plants infected with *ΔnorB*. This heightened defense suggested that bacterially produced NO made the *Rs* cells more visible to plants and could be one reason why the Δ*norB* mutant suffers reduced virulence. We speculate that in addition to protecting itself from oxidative damage, *Rs* may also reduce NO levels in order to hide from its plant hosts. It would be interesting to measure defense responses and bacterial wilt disease susceptibility in plants pretreated with exogenous NO. If high NO levels can alter plant signal transduction, NO-treated plants would have broadly enhanced disease resistance.

## Acknowledgements

The authors thank Corri D. Hamilton and Mark Mandel for helpful discussions and manuscript review.

